# Differential Functional Consequences of *GRIN2A* Mutations Associated with Schizophrenia and Neurodevelopmental Disorders

**DOI:** 10.1101/2023.08.02.551645

**Authors:** Nate Shepard, David Baez-Nieto, Sumaiya Iqbal, Erkin Kurganov, Nikita Budnik, Arthur J. Campbell, Jen Q Pan, Morgan Sheng, Zohreh Farsi

## Abstract

Human genetic studies have revealed rare missense and protein-truncating variants in *GRIN2A*, encoding for the GluN2A subunit of the NMDA receptors, that confer significant risk for schizophrenia (SCZ). Mutations in *GRIN2A* are also associated with epilepsy and developmental delay/intellectual disability (DD/ID). However, it remains enigmatic how alterations to the same protein can result in diverse clinical phenotypes. Here, we performed functional characterization of human GluN1/GluN2A heteromeric NMDA receptors that contain SCZ-linked GluN2A variants, and compared them to NMDA receptors with GluN2A variants associated with epilepsy or DD/ID. Our findings demonstrate that SCZ-associated *GRIN2A* variants were predominantly loss-of-function (LoF), whereas epilepsy and DD/ID-associated variants resulted in both gain- and loss-of-function phenotypes. We additionally show that M653I and S809R, LoF *GRIN2A* variants associated with DD/ID, exert a dominant-negative effect when co-expressed with a wild-type GluN2A, whereas E58Ter and Y698C, SCZ-linked LoF variants, and A727T, an epilepsy-linked LoF variant, do not. These data offer a potential mechanism by which SCZ/epilepsy and DD/ID-linked variants can cause different effects on receptor function and therefore result in divergent pathological outcomes.

## Introduction

NMDA (N-methyl-d-aspartate) receptors (NMDAR) play integral roles in synaptic transmission, development and plasticity^1,2^. Human genetic association studies, as well as postmortem transcriptomics and proteomics studies have implicated synaptic dysfunction in schizophrenia (SCZ) risk^3^. Additionally, several lines of evidence support NMDAR/glutamate hypofunction as a mechanism underlying SCZ pathophysiology: (i) NMDAR antagonists like phencyclidine and ketamine induce SCZ-like symptoms in humans^4^; (ii) autoimmune-NMDAR encephalitis can present with SCZ-like symptoms^5^; (iii) mouse models of NMDAR hypofunction display some phenotypic and biological similarities to SCZ^6^; (iv) extensive human genetics data implicate glutamate receptor signaling alterations in SCZ^7–12^.

The recent Schizophrenia Exome Sequencing Meta-Analysis (SCHEMA) has identified *GRIN2A*, which encodes the GluN2A subunit of the NMDAR, as one of ten genes with exome-wide significance for SCZ risk^13^. *GRIN2A* has additionally been implicated as a SCZ risk gene by genome-wide association studies^14,15^. *GRIN2A* is highly intolerant to mutations leading to a loss-of-function (LoF) phenotype in humans, implying that *GRIN2A* insufficiency is highly detrimental to evolutionary fitness^16^. The association of *GRIN2A* with SCZ is largely driven by protein truncating variants (PTVs)^13^, which are predicted to be loss-of-function (LoF)^17^. A recent multi-omic study of *Grin2a* heterozygous and homozygous null mutant mice has shown that loss of a single copy of *Grin2a* leads to brain-wide transcriptomic changes and phenotypic features reminiscent of human SCZ, including heightened resting gamma oscillation power in electroencephalogram recordings, reduced brain activity in the prefrontal cortex, and a hyperdopaminergic state in the striatum^18,19^.

In addition to SCZ, *GRIN2A* is associated with epilepsy, intellectual disability (ID) and developmental delay (DD)^20^. Association with these neurodevelopmental disorders appears to be mediated predominantly through missense mutations that are often localized in the transmembrane and linker domains of GluN2A^20–22^. Functional analyses of disease-associated variants of *GRIN2A* linked to conditions such as epilepsy or DD/ID have demonstrated both gain- and loss-of-function consequences^22–26^.

Here we present an *in vitro* functional analysis of three groups of *GRIN2A* mutations: i) 11 SCZ-linked mutations found by the SCHEMA study^13^; ii) seven mutations found in the control human subjects of the SCHEMA study^13^ and one mutation from the Genome Aggregation Database (gnomAD)^27^ (referred collectively hereafter as control mutations); iii) four mutations associated with severe DD/ID and epilepsy^20,22,28^. Our data demonstrate that all tested early PTVs and a subset of missense variants associated with SCZ display a LoF phenotype, defined as either reduced current density or an increase in the glutamate EC_50_ or both, while DD/ID-associated missense variants display either a total loss of response to glutamate or a gain-of-function (GoF) effect compared to wild-type receptor function. Additionally, we provide evidence that DD/ID-associated LoF variants can exert a dominant-negative effect when co-expressed with wild-type *GRIN2A*, whereas SCZ-associated LoF variants do not, providing a potential pathomechanistic model that might provide insight into predicting phenotype severity of *GRIN2A* variants.

## Results

### Mutant selection and construct expression in HEK cells

NMDARs are heterotetrameric ligand-gated ion channels that consist of two GluN1 (encoded by *GRIN1*) and two glutamate-binding GluN2 subunits (encoded by *GRIN2A-D*). Each GluN subunit of NMDARs consists of four domains: an extracellular amino-terminal domain (ATD), a clamshell-shaped ligand-binding domain (LBD) which is composed of two non-adjacent segments of the polypeptide, known as S1 and S2, a transmembrane domain (TMD) including three transmembrane helices (M1, M3, and M4) and a membrane re-entrant loop (M2) joined by short linker regions, and an intracellular carboxy-terminal domain (CTD) (Fig. 1A, Fig. S1). We selected a representative set of SCZ-linked mutations found by the SCHEMA study^13^ that reside within different domains of GluN2A including one PTV in the ATD (E58Ter), one PTV (Y700Ter) and four missense mutations in the LBD including two with MPC (Missense badness, PolyPhen-2, Constraint^29^) pathogenicity score > 3 (hereafter referred as mis3; L794M, M788I), and two with 2 < MPC < 3 (hereafter referred as mis2; Y698C, G784A), three missense mutations in the linker/TMD regions (mis3: Q811P; mis2: I605M, G591R), and two mutations in the CTD (mis: I1295T; Frameshift: L1377FS) (Fig. 1A, Fig. S1).

**Figure 1.**
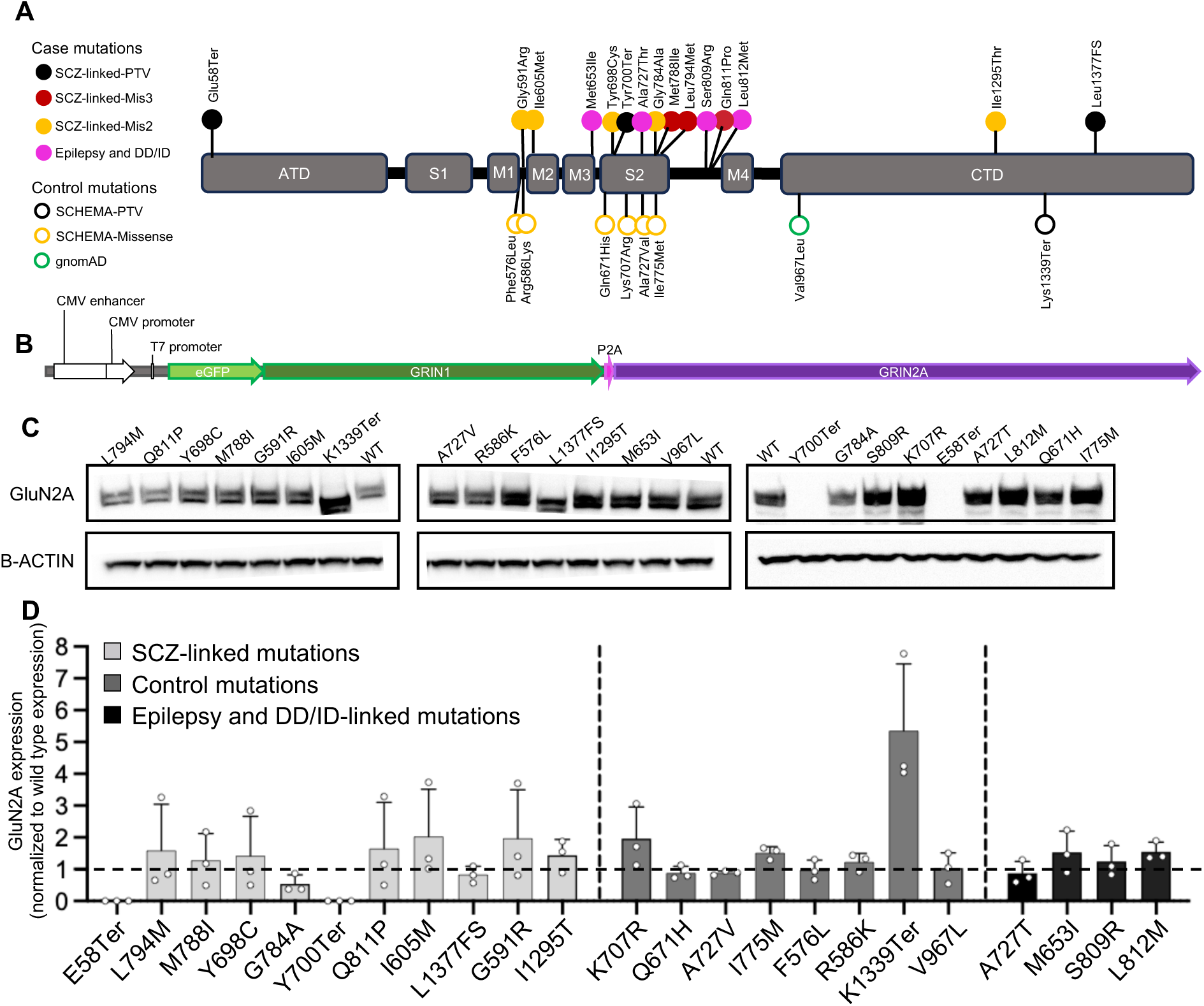
*GRIN2A* mutation selection, construct design, and validation of expression in HEK cells. **(A)** All pathogenic and non-pathogenic variants selected for characterization in this study, mapped on the domain structure of GluN2A. Missense variants from SCZ cases were colored according to the predicted impact (MPC score) in the function. PTV, protein-truncating variant; DD/ID, developmental delay/intellectual disability; FS, frameshift. **(B)** Diagram of the construct transfected into HEK 293T cells for the functional characterization of *GRIN2A* variants. The plasmid was designed with a P2A sequence between the two genes to control the expression of both transcripts with one high efficiency promoter (CMV), and to assure equimolar protein production of GFP-tagged *GRIN1* and wild-type or mutated *GRIN2A* for electrophysiological characterization. **(C)** Western blots probing for GluN2A and β-ACTIN in lysates of HEK cells transiently transfected with *GRIN1-GRIN2A* constructs to express wild-type or mutant NMDARs. The blots presented here are cropped, and the original blots are presented in Supplementary Figure 2. **(D)** Quantification of GluN2A expression by Western blot. All values are normalized to wild-type GluN2A expression. Data are shown as mean + SD; n = 3 for each GluN2A variant. Statistical significance was assessed using Brown-Forsythe and Welch ANOVA with Dunnett’s T3 multiple comparisons test, no conditions were found to be significantly different from wild type.

We also selected eight control mutations: seven of which were identified within a control population of 97,322 individuals from the SCHEMA study^13^. Among the SCHEMA study’s control population, 50,437 individuals had no documented psychiatric diagnoses, and were compiled from 11 global collections that had previously participated in common variant association studies, and 46,885 samples were gathered as part of the gnomAD consortium initiative with no association with psychiatric and neurological conditions. These selected control mutations had no occurrence in SCZ cases and included four in the LBD (mis2: K707R, Q671H, A727V, I775M), two in the M1-M2 linker region (mis2: F576L, R586K), and one in the CTD (PTV: K1339Ter). We also selected one missense mutation from the gnomAD database in the CTD, V967L, because it is the most prevalent *GRIN2A* missense variant in the general population and has no clinical phenotype (Fig. 1A, Fig. S1).

Additionally, to elucidate the specific effects of SCZ-linked mutations, we characterized one epilepsy-associated mutation A727T^28^ in the LBD, and three missense *GRIN2A* mutations associated with DD/ID in the linker/TMD regions: M653I, S809R, and L812M^20,30^ the latter of which has previously been characterized^22^ (Fig. 1A, Fig. S1).

We transfected HEK cells with plasmids encoding for GFP-tagged human *GRIN1* fused with a self-cleaving 2A peptide and wild-type or mutated human *GRIN2A*, allowing us to express GluN1 and GluN2A proteins at a 1:1 ratio (Fig. 1B). We confirmed robust expression of the constructs after 24h visually as the majority of cells were GFP-positive. Biochemistry analysis by Western blot, showed that cells transfected with PTVs located in the N-terminal half of the protein, E58Ter and Y700Ter, showed no detectable expression of GluN2A at the size corresponding to wild-type GluN2A (Fig. 1C, Fig. S2). In case of Y700Ter, a faint band near the expected size for the Y700Ter fragment (∼ 80 kDa) was observed (Fig. S2). Both E58Ter and Y700Ter introduce premature termination codons in their mRNA transcripts, leading to their degradation by nonsense-mediated decay (NMD). NMD is a well-established cellular mechanism designed to degrade mRNA transcripts containing premature termination codons or nonsense mutations^17^. However, PTV transcripts are more likely to escape NMD when they are downstream of 50-55 nucleotides before the last exon-exon junction^31^, or reside in the last exon^32^. This could explain why the PTVs nearer to the carboxy-terminal end of the protein (L1377Frameshift and K1339Ter, both located in the last exon of *GRIN2A*) did not lead to decreased protein expression in HEK cells. For the K1339Ter variant, an approximately fivefold increase in expression was observed compared to the wild-type GluN2A (p = 0.07; Fig. 1D). The increased expression could potentially be attributed to the functional disruption of a di-leucine motif positioned at residues 1319-1320, situated in close proximity to the K1339Ter mutation site. This motif has been established to play a regulatory role in the endocytosis of NMDARs containing GluN2A^33^. All *GRIN2A* missense variants tested, except G784A, were expressed at comparable or higher levels than wild-type GluN2A (Fig. 1C-D).

### Functional characterization of SCHEMA and non-SCHEMA mutations

We next recorded whole-cell currents from HEK cells expressing wild-type or mutant NMDARs using a high-throughput automated planar patch-clamp system (Syncropatch 384PE). Cells were held at -60 mV during the recording and currents were evoked using the liquid handler to “puff” increasing concentrations of glutamate (1, 3, 10, 30, and 100 μM) in the constant presence of a saturating concentration of glycine (30 μM) (Fig. 2A). Of the 11 selected SCZ-linked variants, both PTVs located in the N-terminal half of the protein (E58Ter, Y700Ter) as well as one mis3 (Q811P) and one mis2 (Y698C) variants displayed electrophysiological characteristics that were significantly different from the wild-type control (Fig. 2B-C). E58Ter and Y700Ter showed no response to any concentration of glutamate (Fig. 2B), consistent with the lack of protein expression of these mutants (Fig. 1C-D). The missense variant Q811P, predicted to have a high degree of deleteriousness (mis3), showed a fivefold increase in glutamate EC_50_ compared to the wild-type control (Q811P: 19.91 ± 2.11 µM; WT: 3.92 ± 0.257 µM, p < 0.0001) (Fig. 2C), but no change in maximal response at a saturating concentration (100 µM) of glutamate. The mis2 variant Y698C showed a fivefold reduction in maximal response to the saturating concentration of glutamate compared to wild type (Y698C: -10.40 ± 1.47 pA/pF; WT: -46.88 ± 8.67 pA/pF, p < 0.0001) (Fig. 2B) as well as a twofold increase in glutamate EC_50_ (Y698C: 6.55 ± 0.731 µM; WT: 2.79 ± 0.255 µM, p < 0.05) (Fig. 2C). The other tested SCZ-linked mutations did not significantly differ in any of the measured characteristics compared to the wild-type GluN2A/GluN1 NMDARs (Fig. 2D-E, Fig. S3; Table 1).

**Figure 2.**
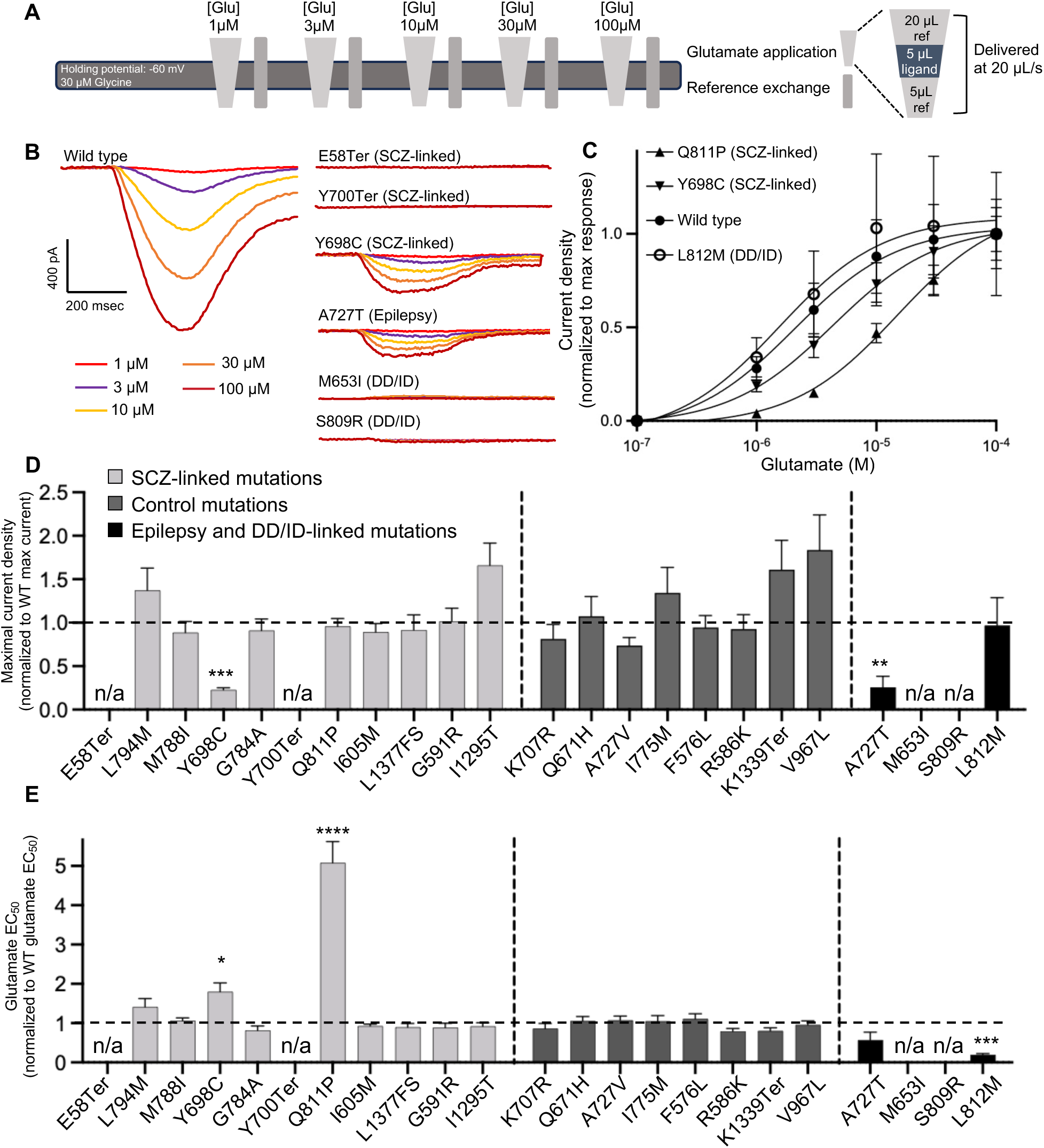
Schizophrenia and DD/ID-associated *GRIN2A* mutations demonstrate both gain- and loss-of-function effects. **(A)** Whole-cell recording protocol using Syncropatch to record NMDAR currents from HEK cells transiently transfected with the construct shown in Fig.1B. Cells were puffed with different concentrations of glutamate stacked in between different volumes of reference solution (light gray triangles, inset on the right panel). Each puff results in a ∼250 ms transient of glutamate exposure to the receptors. After each glutamate application half of the volume of the well was replaced with fresh reference solution (“Reference exchange”, gray bars) to minimize desensitization due to remnant glutamate in the well. Glu, glutamate; ref, reference solution. **(B)** Averaged current traces of wild-type and selected mutant NMDARs evoked by ∼250 ms transients of glutamate exposure of increasing concentration, in the presence of 30 μM glycine. The traces were colored according to the different concentrations of glutamate. **(C)** Averaged current density in response to increasing concentrations of glutamate in the constant presence of 30 μM glycine, normalized to maximal response, for wild-type and selected mutant NMDARs. The lines indicate a nonlinear regression three-parameter fit to each dataset. **(D)** Peak current density in response to 100 µM glutamate, normalized to wild type’s response, for each mutant NMDAR. (E) Glutamate EC_50_ normalized to wild type EC_50_, for each mutant NMDAR. In **C-E** data are displayed as mean ± SEM, n = 10-77; see Table 1 for number of cells recorded per variant; statistical significance was assessed using Brown-Forsythe and Welch ANOVA with Dunnett’s T3 multiple comparisons test. *: p < 0.05, ***: p < 0.001, ****: p < 0.0001.

**Table 1.** Electrophysiological and expression data for all characterized *GRIN2A* mutations. Data are given as mean +/-SEM except for protein expression values which are given as mean +/-SD. Statistical significance was assessed using Brown-Forsythe and Welch ANOVA with Dunnett’s T3 multiple comparisons test. Bolded text indicates significantly different values from wild type. *: p < 0.05, **: p < 0.01, ***: p < 0.001, ****: p < 0.0001. The cell colors in the table indicate the pathological status of the variants. The lightest gray shade represents the SCZ-linked mutation, the second light gray corresponds to control mutations, and the darkest gray shade is indicative of mutations linked to epilepsy and developmental delay/intellectual disability (DD/ID).

| Variant | Location on GluN2A | Mutation Class | Glutamate EC<sub>50</sub> (normalized to WT) | Max Amplitude (normalized to WT) | Protein Expression (normalized to WT) | Number of Cells |
| --- | --- | --- | --- | --- | --- | --- |
| E58Ter | ATD | PTV | N/A | N/A | N/A | 41 |
| L794M | LBD (S2) | mis3 | 1.41 ± 0.21 | 1.37 ± 0.26 | 1.59 ± 0.84 | 37 |
| M788I | LBD (S2) | mis3 | 1.06 ± 0.08 | 0.89 ± 0.13 | 1.28 ± 0.84 | 71 |
| G784A | LBD (S2) | mis2 | 0.82 ± 0.11 | 0.91 ± 0.13 | 0.54 ± 0.16 | 33 |
| Y698C | LBD (S2) | mis2 | **1.80 ± 0.23 (\*)** | **0.23 ± 0.02 (\*\*\*\*)** | 1.42 ± 0.72 | 47 |
| Y700Ter | LBD (S2) | PTV | N/A | N/A | N/A | 36 |
| Q811P | Linker (S2-M4) | mis3 | **5.08 ± 0.54 (\*\*\*\*)** | 0.96 ± 0.09 | 1.65 ± 0.84 | 50 |
| I605M | TMD (M2) | mis2 | 0.93 ± 0.05 | 0.89 ± 0.10 | 2.03 ± 0.86 | 81 |
| L1377FS | CTD | PTV | 0.90 ± 0.09 | 0.92 ± 0.18 | 0.83 ± 0.15 | 31 |
| G591R | Linker (M1-M2) | mis2 | 0.89 ± 0.11 | 1.01 ± 0.15 | 1.97 ± 0.88 | 58 |
| I1295T | CTD | mis2 | 0.92 ± 0.09 | 1.66 ± 0.25 | 1.44 ± 0.29 | 37 |
| K707R | LBD (S2) | mis2 | 0.87 ± 0.13 | 0.81 ± 0.17 | 1.97 ± 0.57 | 18 |
| Q671H | LBD (S2) | mis2 | 1.06 ± 0.11 | 1.07 ± 0.23 | 0.88 ± 0.12 | 17 |
| A727V | LBD (S2) | mis2 | 1.07 ± 0.11 | 0.74 ± 0.09 | 0.88 ± 0.04 | 70 |
| I775M | LBD (S2) | mis2 | 1.05 ± 0.14 | 1.35 ± 0.29 | 1.51 ± 0.11 | 36 |
| F576L | Linker (M1-M2) | mis2 | 1.11 ± 0.13 | 0.95 ± 0.14 | 0.97 ± 0.18 | 35 |
| R586K | Linker (M1-M2) | mis2 | 0.79 ± 0.07 | 0.93 ± 0.17 | 1.23 ± 0.16 | 38 |
| V967L | CTD | mis2 | 0.81 ± 0.07 | 1.84 ± 0.40 | 1.03 ± 0.28 | 38 |
| K1339Ter | CTD | PTV | 0.96 ± 0.10 | 1.61 ± 0.34 | 5.35 ± 1.21 | 33 |
| A727T | LBD (S2) | mis2 | 0.57 ± 0.20 | **0.26 ± 0.13 (\*\*)** | 0.89 ± 0.21 | 10 |
| L812M | Linker (S2-M4) | mis3 | **0.20 ± 0.03 (\*\*\*)** | 0.97 ± 0.32 | 1.55 ± 0.17 | 13 |
| S809R | Linker (S2-M4) | mis3 | N/A | N/A | 1.24 ± 0.29 | 30 |
| M653I | TMD (M3) | mis3 | N/A | N/A | 1.53 ± 0.38 | 77 |

SCZ-linked mutations
Control mutations

Together, our data demonstrate that SCZ-linked missense *GRIN2A* variants either had no measurable effect on NMDAR function in our electrophysiological assay or resulted in reduced glutamate potency or efficacy. Notably, the Q811P and Y698C variants demonstrated a LoF phenotype, similar in direction but less severe than the SCZ-linked PTVs. Consistent with our results, prior characterization of another SCZ-linked missense variant identified by the SCHEMA study, A716T, revealed a LoF phenotype of increased glutamate EC_50_^25^.

None of the control variants tested (seven from the SCHEMA study and one from gnomAD) showed a significantly different EC_50_ or maximal response to glutamate (Fig. 2D-E, Fig. S4; Table 1).

All four epilepsy and DD/ID-linked mutations showed significantly different electrophysiological phenotypes from wild type (Fig. 2B-E, Fig. S5). A727T, associated with epilepsy^28^, showed a significant reduction in maximal response to glutamate (A727T: -6.595 ± 3.13 pA/pF; WT: -29.01 ± 4.54 pA/pF, p < 0.005) (Fig. 2B, D), consistent with previous characterizations^25^. L812M displayed a fivefold reduction in glutamate EC_50_ (L812M: 1.90 ± 0.266 µM; WT: 9.66 ± 1.53 µM, p < 0.0001) (Fig. 2C, E) and no change in maximal current density (Fig. 2D), consistent with previous studies showing that this variant enhances glutamate potency^22^. By contrast, both M653I and S809R did not respond to any concentration of glutamate, despite protein expression levels similar to wild type (Fig 1D), indicating a LoF phenotype (Fig. 2B, Fig. S5).

In summary, out of the 11 SCZ-linked *GRIN2A* variants characterized by this study, two tested PTVs located in the N-terminal half of the protein led to complete loss of current in response to glutamate and two out of the eight missense variants showed a partial LoF phenotype. Of the eight characterized control variants, seven missense variants and one PTV, none demonstrated any changes as compared to the wild-type receptor. Notably, all tested epilepsy and DD/ID-associated missense variants were significantly different from wild type and displayed either a severe LoF or a GoF phenotype (Fig. 2 D-E; Table 1).

### Dominant-negative phenotype as a differentiator of severe DD/ID and epilepsy/SCZ

Our data demonstrate that *GRIN2A* missense variants associated with either epilepsy, DD/ID or SCZ can result in LoF of the NMDAR. However, the clinical features of these disorders, all of which stem from heterozygous mutation of *GRIN2A* in humans, are distinct, raising the question whether there are functional differences in the *in vivo* consequences of these variants. In order to better reflect the human disease state, we co-expressed the SCZ, epilepsy, and DD/ID-linked variants, that were found to result in LoF, with the wild-type variant and investigated their effects on NMDAR function. Having previously demonstrated that the DD/ID-linked missense variant M653I produces a comparable amount of protein as the wild-type receptor (Fig. 1C-D), we first tested whether the null electrophysiological phenotype caused by this variant in heterologous cells was a result of dysfunctional trafficking of the receptor to the cell surface. To this end, we conducted surface biotinylation of HEK cells expressing either the M653I variant or wild-type GluN2A. We validated the assay by also probing for insulin receptor B, endogenously expressed on the surface of HEK cells, and beta-actin, endogenously expressed primarily in the cytoplasm. This assay demonstrated that M653I-containing NMDARs are present at the cell surface at a comparable level to the wild-type receptor (Fig. 3A, Fig. S6).

**Figure 3.**
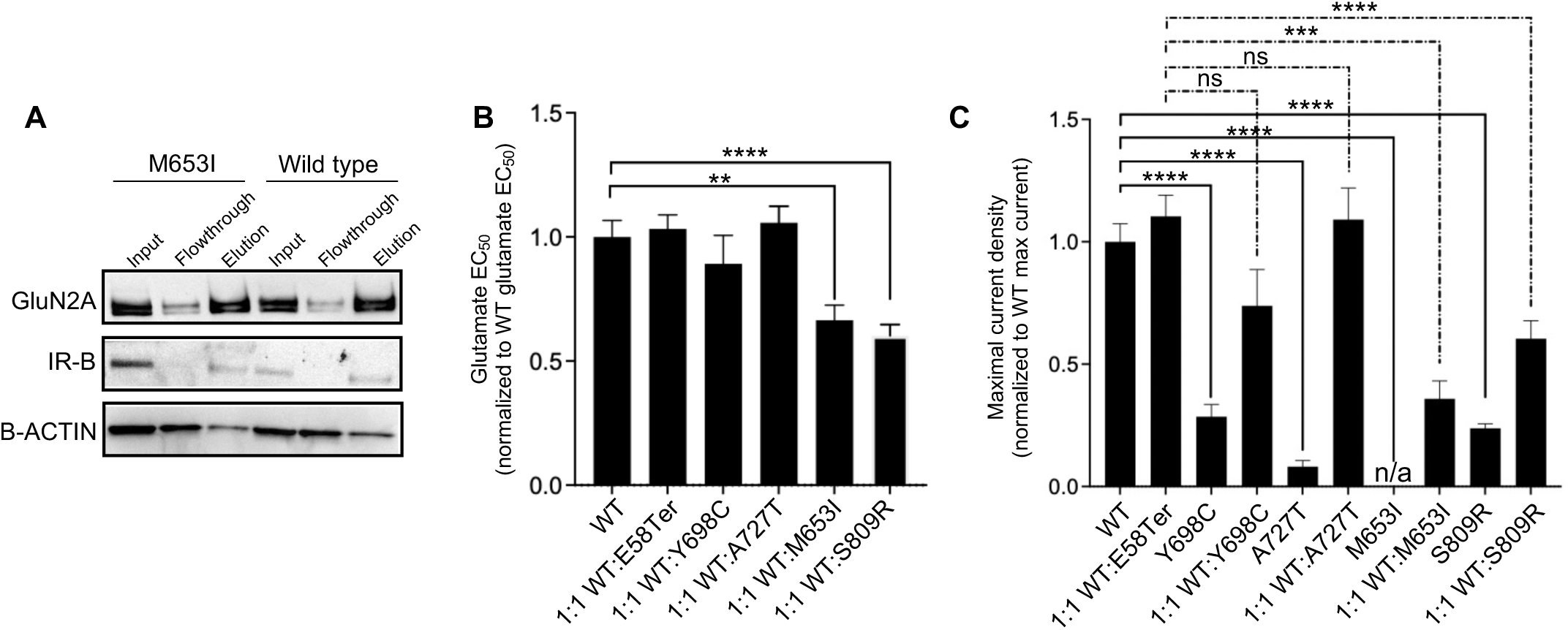
*GRIN2A* mutations associated with DD/ID, but not epilepsy or SCZ, demonstrate a dominant-negative effect. **(A)** Western blot probing for GluN2A, insulin receptor beta, and β-ACTIN in the input, flowthrough, and elution samples of surface biotinylation experiment done on HEK cells transiently transfected with *GRIN1-GRIN2A* constructs to express wild-type or M653I mutant NMDARs. Input, flowthrough, and elution represent total, internal, and surface expression respectively. IR-B: insulin receptor beta. The blots presented here are cropped, and the original blots are presented in Supplementary Figure 6. **(B)** Glutamate EC_50_ normalized to wild type EC_50_, for each mutant NMDAR. **(C)** Peak current density in response to 100 µM glutamate, normalized to wild type’s response, is plotted for each mutant NMDAR. In **B** and **C** data are displayed as mean +/-SEM; n = 80 (WT); 113 (1:1 WT:E58Ter); 27 (Y698C); 45 (1:1 Y698C:WT); 8 (A727T); 44 (1:1 A727T:WT); 74 (M653I); 39 (1:1 WT:M653I); 35 (S809R); 46 (1:1 WT:S809R) cells; statistical significance was assessed using Brown-Forsythe and Welch ANOVA with Dunnett’s T3 multiple comparisons test. *: p < 0.05, ***: p < 0.001, ****: p < 0.0001.

We then tested whether either SCZ-linked or epilepsy and DD/ID-linked variants would exert a dominant-negative effect on NMDAR function when co-expressed with an equal amount of wild-type *GRIN2A* (as well as the obligate *GRIN1* subunit). For this experiment, we included the DD/ID-linked variants, M653I and S809R, the epilepsy-linked variant, A727T, as well as Y698C, a SCZ-linked missense variant, all with a LoF phenotype (Fig. 2B-E), and E58Ter, an early termination mutation that produces no protein (Fig. 1C) and therefore should not have a dominant-negative phenotype. No difference in glutamate EC_50_ or maximal current density was observed between the 1:1 WT:E58Ter co-expressing cells and the wild-type NMDAR (Fig. 3B-C). Interestingly, the 1:1 WT:Y698C and 1:1 WT:A727T co-expressing cells demonstrated no change in glutamate EC_50_ and their maximal responses did not exhibit a statistically significant difference when compared to cells co-expressing 1:1 WT:E58Ter (Fig. 3B-C). Cells co-expressing 1:1 WT:M653I and 1:1 WT:S809R *GRIN2A*, however, showed a decrease in glutamate EC_50_ and a significant reduction of ∼70**%** and ∼45% in maximal current compared with cells co-expressing 1:1 WT:E58Ter, respectively (WT:M653I vs WT:E58Ter: p < 0.0001; WT:S809R vs WT:E58Ter: p = 0.0002; Fig. 3B-C), demonstrating that the DD/ID-linked variants, M653I and S809R, exert a dominant-negative effect on NMDAR function, whereas the epilepsy-linked variant, A727T, and the SCZ-linked variants, Y698C and E58Ter, do not.

These data uncover a potential mechanism by which similar functional consequences (LoF) of GluN2A alteration can lead to different effects on receptor function, and therefore divergent pathological outcomes.

## Discussion

A growing number of disease-associated variants have been identified in *GRIN2A*, highlighting the need for mechanistic data to establish connections between genetic variants and the resultant pathological phenotypes. Here, we present electrophysiological data for a representative set of *GRIN2A* variants associated with SCZ, epilepsy or DD/ID, and propose a pathomechanistic model that could potentially aid in predicting phenotype severity of *GRIN2A* variants.

In agreement with previous findings in *GRIN2A*, *GRIN1* and *GRIN2B*^20,25,34,35^, our data suggest that the location of mutations in the structure of GluN2A can inform predictions of their functional consequences and pathogenicity. We noted that all disease-associated *GRIN2A* missense variants that led to an electrophysiological phenotype significantly different from wild type were located either within the LBD (Y698C and A727T) or linker/TMD regions (Q811P, L812M, S809R and M653I) of GluN2A while none of the mutations putatively localized in the CTD (I1295T, L1377FS, V967L, and K1339Ter) demonstrated electrophysiological changes. Additionally, we show that among disease-associated *GRIN2A* variants, the DD/ID mutations localized within the linker/TMD region of GluN2A resulted in either a severe LoF (no current) or GoF (enhanced glutamate potency) consistent with prior characterizations of mutations from this regions^20,22–25,28,36^, while SCZ/epilepsy-linked missense mutations in the LBD displayed either no effect or partial LoF (reduced current or glutamate potency). These data align with previous studies showing that missense variants in the linker/TMD regions are associated with more severe DD/ID, while missense variants in the LBD generally result in LoF and are associated with less severe abnormalities with mild to no ID^20,25^.

It is noteworthy, however, that heterologous overexpression of *GRIN2A* variants in HEK cells may not reveal deficits in trafficking and other post-translational mechanisms resulting from these mutations. For instance, reduced glutamate potency has been shown to often be associated with decreased surface expression of NMDARs in neurons^25^. Thus, it is possible that disease-associated missense mutations that lead to reduced glutamate potency in our study (such as Q811P and Y698C) lead to a more severe LoF phenotype when expressed in neurons due to cumulative effects on glutamate potency and receptor trafficking. Additionally, it has been shown that rare variants at the interfaces between subunits of NMDARs might not necessarily affect the receptor function but lead to impaired receptor trafficking^25^ resulting in a LoF phenotype. An example of such a variant in our study is the SCZ-linked L794M which forms a non-bonded contact with GluN1 upon NMDAR assembly. Although we did not observe any effect of L794M on NMDAR function, it is plausible that its expression in neurons results in LoF. Finally, the heterologous overexpression system might not reveal the functional consequences of mutations in the CTD which has an indispensable role in NMDAR-mediated intracellular signaling and synaptic physiology. Thus, further studies of the tested *GRIN2A* variants expressed in neurons are necessary to elucidate the impact of these mutations in a physiologically relevant context.

Our data also offers a potential mechanistic explanation for how missense *GRIN2A* variants can result in the same phenotype in the homozygous state but lead to distinct neurological manifestations in a heterozygous state. We show that the DD/ID-linked M653I and S809R, epilepsy-linked A727T and SCZ-linked Y698C *GRIN2A* variants all result in LoF, but, when co-expressed with wild-type *GRIN2A*, only M653I and S809R exert a dominant negative effect. Looking into the position of these mutations in the 3D structure of the GluN1/GluN2A NMDAR, we noted that the M653I and S809R mutations, but not Y698C or A727T, are located at the interface between GluN2A and GluN1, and introduce intramolecular interactions with GluN1, which are not present in the wild-type NMDAR. Thus, it is possible that the new interactions introduced by M653I and S809R mutations are rigidifying the assembly and thereby causing a negative effect on the entire assembly. This is consistent with previous findings showing that mutations in the protein-protein interface can cause a dominant negative effect by poisoning the assembly of the protein complex^37,38^. These data suggest that the consequences of *GRIN2A* haploinsufficiency as a result of DD/ID-associated missense variants might be more severe as compared to SCZ/epilepsy-associated missense variants and PTVs, despite the two groups having similar effects on NMDAR function in a homozygous state. These data highlight the importance of analyses of *GRIN2A* variants in both homozygous and heterozygous conditions to effectively differentiate between various potential mechanisms. Together, our findings offer a better understanding of the relationship between genetic variant, NMDAR dysfunction, and disease phenotype, and could potentially contribute to development of pharmacologic strategies to correct NMDAR function.

## Methods

### DNA Constructs

The wild-type construct was synthesized by Genscript Biotech by adding eGPF (accession JN204884.1), GRIN1 (accession NM_000832.7), and GRIN2A (accession NM_000833.5) to the pcDNA3.1+P2a backbone. Mutant constructs were generated by Genscript Biotech using site-directed mutagenesis. All constructs were transformed into NEB Turbo Competent E. coli (High Efficiency) (New England Biolabs C2984H) and purified using the NucleoBond Xtra Maxi EF kit (Machery Nagel 740424).

### Cell Culture and Electroporation

HEK 293-T cells (Sigma-Aldrich 12022001) were cultured at 37 °C, 5% CO2 in DMEM, high glucose, GlutaMAX Supplement, pyruvate (Thermo 10569044) supplemented with 10% fetal bovine serum and 5% pen-strep (DMEM + FBS/PS). Cells were electroporated using the MaxCyte STX electroporator. Briefly, 50-75% confluent cells were washed with PBS, lifted, pelleted, and resuspended at a final density of 100 million cells/mL in electroporation buffer (MaxCyte proprietary buffer). 100 µL of this mixture was loaded in the electroporation cassette (OC-100), and electroporated using the preset “HEK 293” protocol with 25 µg of the appropriate DNA construct concentrated to ≥ 5 µg/µL. For co-expression in the dominant-negative experiments, cells were electroporated with a mixture of 12.5 µg of both constructs. Post-transfection, 10 mg of DNAse I (Stemcell Technologies 07900) was added to the cells, and they were allowed to recover for 30 min at 37 °C, 5% CO2, and then rescued into DMEM + FBS/PS containing 15 µM DCKA, 8 µM APV, and 2 µM MK801.

### Whole-Cell Recording

Cells were recorded using the Syncropatch 384PE 24h after electroporation in a whole-cell configuration. Transfected cells were washed twice with PBS and treated with Accutase (Stemcell Technologies 07920) for 5 min, and then pelleted and resuspended in extracellular solution (in 8 mM glucose, 4 mM KCl, 10 mM HEPES, and 145 mM NaCl (pH 7.4)) at a density of 400K cells/mL. We used a cesium-containing internal recording solution to minimize the contribution of endogenous cationic current during the recording (in 20 mM EGTA, 10 mM CsCl, 110 mM CsF, 10 mM HEPES, and 10 mM NaCl, pH 7.2).

The patch process consisted of a series of different negative pressures and negative voltages to foster the giga-seal to the glass (REF PMID: 29736723). To enhance the seal formation the cells were transiently exposed to a high Ca^2+^ recording solution “seal enhancer solution” (in 80 mM NaCl, 8 mM glucose, 60 mM NMDG, 4 mM KCL, 10 mM HEPES, and 10 mM CaCl_2_, pH 7.4). The cells were washed four times (replacing half of the volume of the well) with standard recording solution (in mM 80 NaCl, 8 glucose, 60 NMDG, 4 KCL, 10 HEPES, and 6 CaCl_2_, and 30 μM glycine, pH 7.4) before starting the recording. After gaining electrical access (whole-cell configuration), cells were held at a holding potential of -60 mV, and NMDAR currents were evoked by a “puffing addition protocol” (PMID: 36340694) with increasing concentrations of glutamate (0, 1, 3, 10, 30, 100 µM) in recording solution. Glutamate puffs exposed the cells to the glutamate-containing solution for ∼250 ms before being washed out with the recording solution in the stack. All the glutamate was removed completely after each puff. To remove any traces of glutamate, after each puff half of the volume of the well was replaced with fresh recording solution in between each addition of ligand.

### Electrophysiological analyses

Groups of 5 variants at a time were transfected and recorded along with a wild-type construct as a reference, and the results for each of the variants were normalized to this wild-type control. Different batches of cells electroporated with wild-type control showed a variability in the maximal response to glutamate, but not in the glutamate EC_50_, likely related to the electroporation efficiency for each particular batch of cells. Cells were selected to be analyzed based on having a seal resistance of ≥ 50 MOhm for all ligand application steps and a maximal current ≥ 50 pA. Traces were analyzed between 800 ms and 1400 ms after initiation of stacked ligand addition. All current values were normalized to the capacitance of the recorded cell to calculate current density. Glutamate EC_50_ values were calculated for each cell using a 3-parameter agonist-response model with a Hill slope of 1.0, Response = Bottom + Concentration*(Top-Bottom)/(EC50 + Concentration).

### Immunoblot analysis

Cells were electroporated as described. After 24h, cells were washed 2x with PBS and lysed in 1% SDS containing protease inhibitors (Sigma 4693159001) and nuclease (Sigma E1014). The protein concentrations of cell lysate were determined using Bicinchoninic acid assay (BCA; Pierce 23227). To equalize protein concentrations, samples were diluted with 4X SDS-Sample buffer (Boston Bioproducts BP-111R; to a final concentration of 1X) and water. The diluted samples were then left at room temperature for 20 min. 25 µg of protein in an equal volume were loaded for each sample on 3-8% Tris-acetate polyacrylamide gels; the gels were run using Tris-acetate SDS running buffer at constant voltage. Proteins were transferred to 0.2 μm Nitrocellulose membranes using semi-dry transfer (BioRad Transblot Turbo; 25V 30 min). Membranes were blocked using 5% milk in Tris-buffered saline supplemented with 0.1% Tween-20 (TBST) for 1.5h at room-temperature (RT). Membranes were then probed overnight at 4°C with primary antibody (Novus rabbit anti-NMDAR2A NB300-105, 1:2000 or CST rabbit anti-Insulin Receptor β 4B8, 1:1000) in 1% milk TBST with gentle rotation. After three 10-min washes in TBST, membranes were incubated with 1:5,000 HRP-conjugated (Jackson Immunoresearch) anti-IgG antibody in 1% milk TBST for 60 min at RT. All membranes were then washed three times in TBST and imaged on the ChemiDoc MP (BioRad) platforms. Membranes were then re-probed with an HRP-conjugated mouse anti-beta actin antibody (Sigma A3854) for 1h at RT in 5% milk TBST and imaged again as described.

### Surface Biotinylation

Cells were electroporated as described before. After 24h, cells were washed two times with PBS supplemented with 1 mM CaCl_2_ and 1 mM MgCl_2_. Cells were then incubated in 1 mg/mL sulfo-NHS-LC biotin in PBS + Mg/Ca (Thermo 21335) on a flat surface for 30 min-2h at 4 °C. Cells were then quenched in 20 mM Tris in PBS + Mg/Ca for 15 min at 4°C. Cells were then lysed in 20 mM Tris.Cl (pH 7.5), 150 mM NaCl, 0.2% Triton-X 100, 1% SDS in water supplemented with protease inhibitors and nuclease. After quantification of lysate with BCA, 100 µg of lysate was removed and diluted to 0.2% SDS by water, and rotated at 4 °C for 30 min. 80 µL of neutravidin-agarose beads (Thermo 29204) was added to this lysate and the solution was rotated overnight at 4 °C. Afterwards, the solution was spun down, and a sample of the flowthrough was taken for analysis, and the remaining beads were washed in 200 µL lysis buffer three times. Beads were then eluted with 4X SDS-Sample buffer.

### Statistics

All quantitative data is given as mean ± SEM, unless stated differently. All statistical comparisons were done using one-way Brown-Forsythe and Welch ANOVAs with Dunnett’s T3 multiple comparison correction.

## Acknowledgements

We thank Andrew Allen and Ned Martins for assistance with the construct design and Syncropatch measurements. We also thank Sameer Aryal, Bryan Song and Michel Weiwer for invaluable feedback. Research reported in this manuscript was supported by the Stanley Center for Psychiatric Research.

## Author contributions

Z.F. and M.S. designed the study with assistance from D.B.N. and J.Q.P.. N.S., N.B. and E.K. performed all the experiments with assistance from D.B.N.. N.S., D.B.N., and Z.F. performed data analysis. S.I. and A.J.C. carried out the protein structure analysis. N.S. and Z.F. wrote the manuscript with inputs from all co-authors.

## Data availability statement

All data generated or analyzed during this study are included in this published article (and its Supplementary Information files), and are available from the corresponding author on reasonable request.

## Additional Information

M.S. is cofounder and SAB member of Neumora Therapeutics, and serves on the SAB of Biogen, ArcLight, Vanqua Bio, and Proximity Therapeutics.

## Supplemental Figure Legends

**Figure S1.**
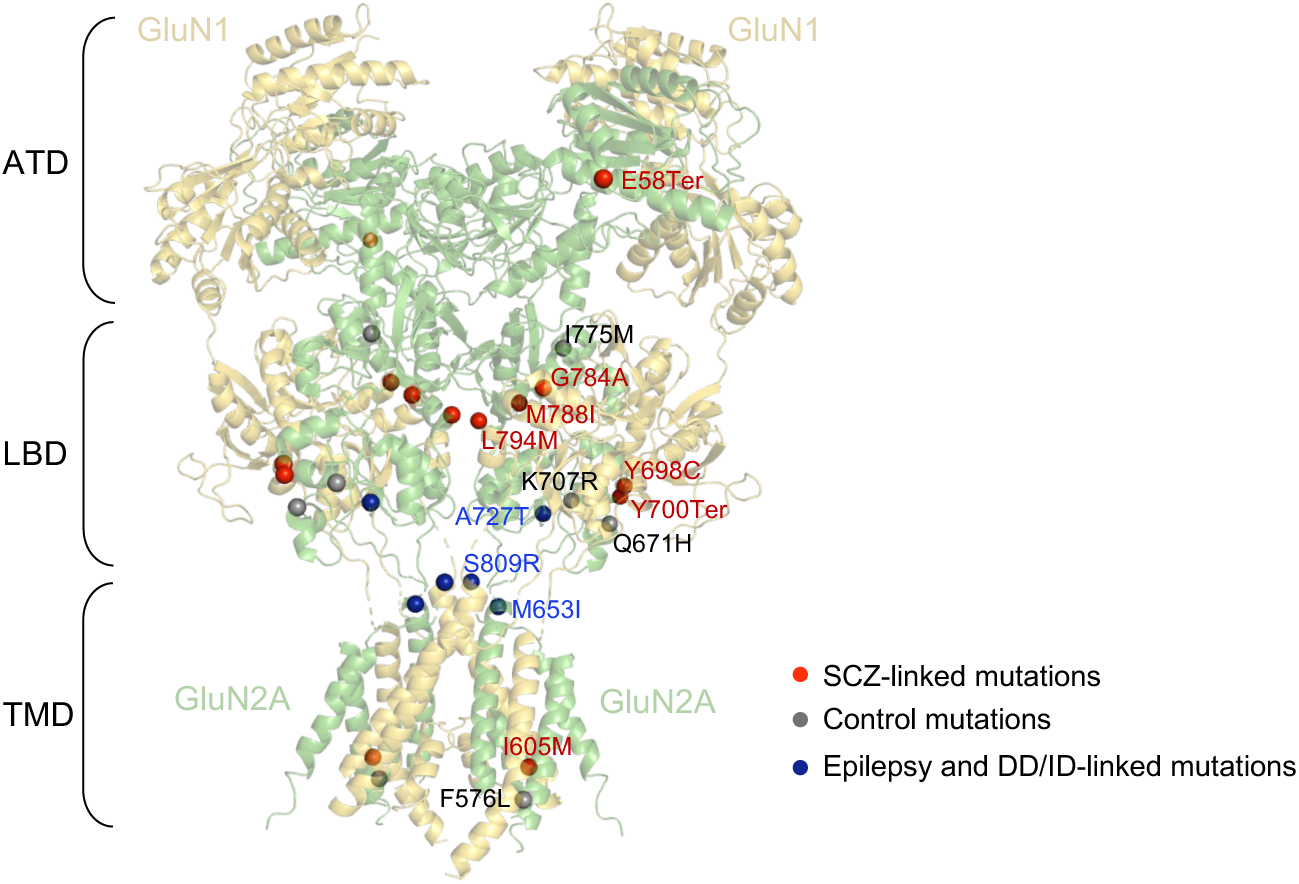
*GRIN2A* mutations mapped on human NMDAR structure. Protein structure of human NMDAR (PDB ID 6IRH) with two GluN1 (light yellow) and two GluN2A (light green) subunits. *GRIN2A* mutations associated with SCZ (red) or epilepsy/DD/ID (blue) as well as control mutations (dark gray) are mapped on the two GluN2A subunits of human NMDAR. Two SCZ-associated missense mutations (Q811P, G591R), one control mutation (R586K), one DD/ID-linked mutation (L812M) as well as mutations within the CTD were not mapped due to no structure coverage.

**Figure S2.**
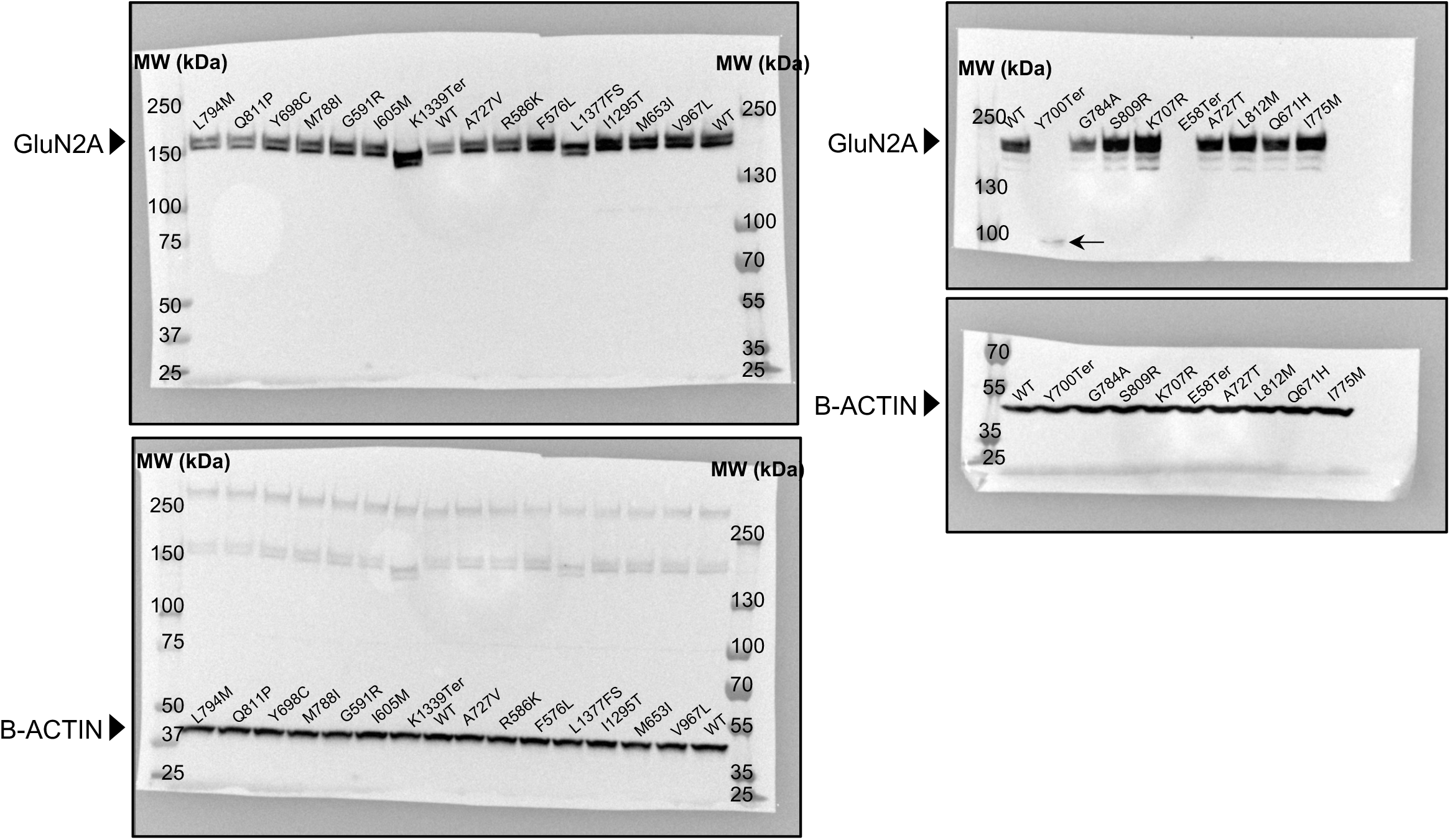
Uncropped western blot images of GluN2A variants. Western blots probing for GluN2A and β-ACTIN in lysates of HEK cells transiently transfected with GRIN1-GRIN2A constructs to express wild-type or mutant NMDARs. The arrow in Y700Ter lane indicates a faint band near the expected size for the Y700Ter fragment (∼ 80 kDa).

**Figure S3.**
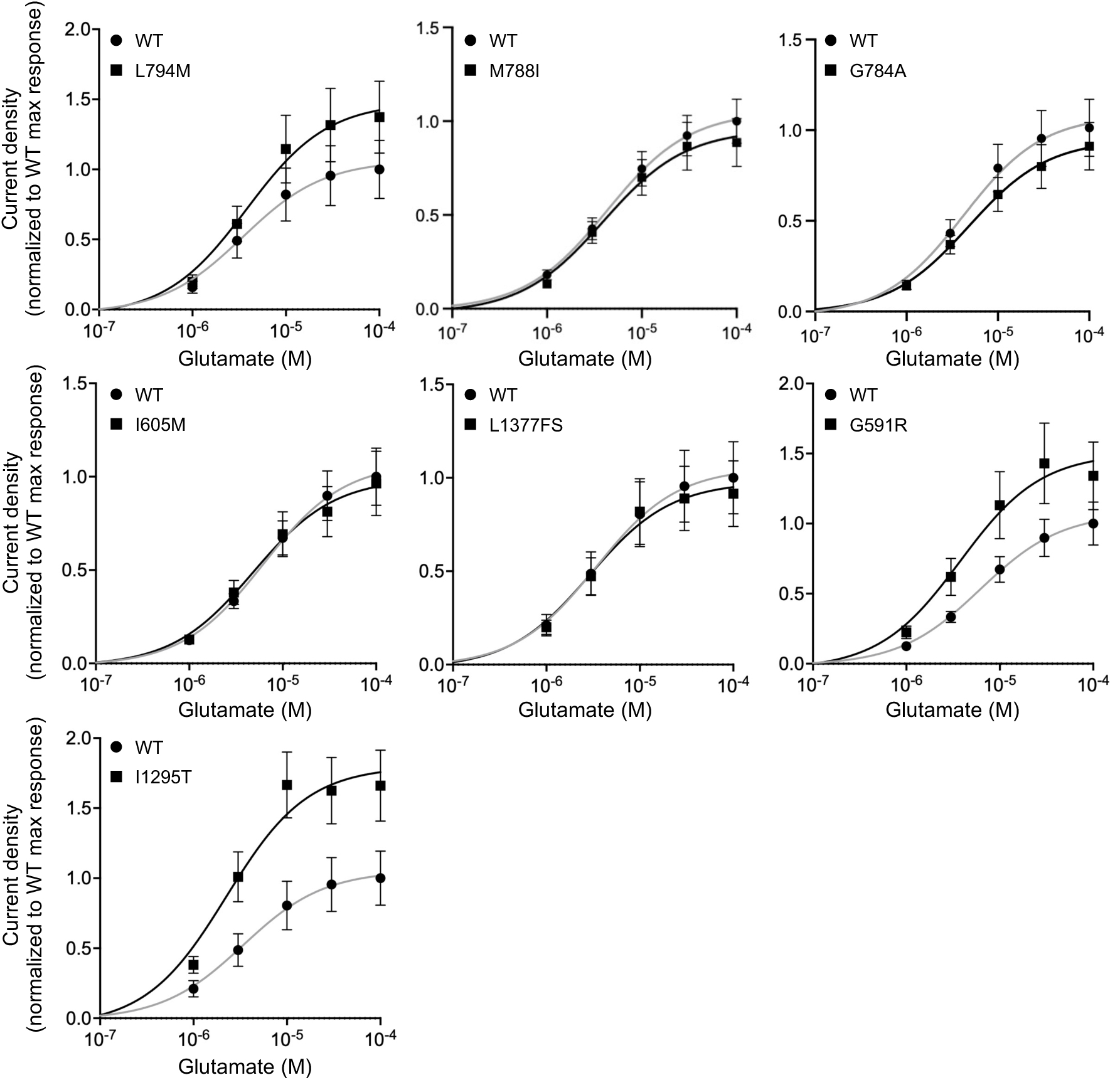
Effect of SCZ-associated *GRIN2A* variants on NMDAR function. Averaged current density in response to increasing concentrations of glutamate in the constant presence of 30 μM glycine, normalized to maximal response, for NMDARs containing the wild-type or SCZ-associated GluN2A variants. The lines indicate a nonlinear regression three-parameter fit to each dataset. Data are displayed as mean ± SEM, n = 31-81; see Table 1 for number of cells recorded per variant.

**Figure S4.**
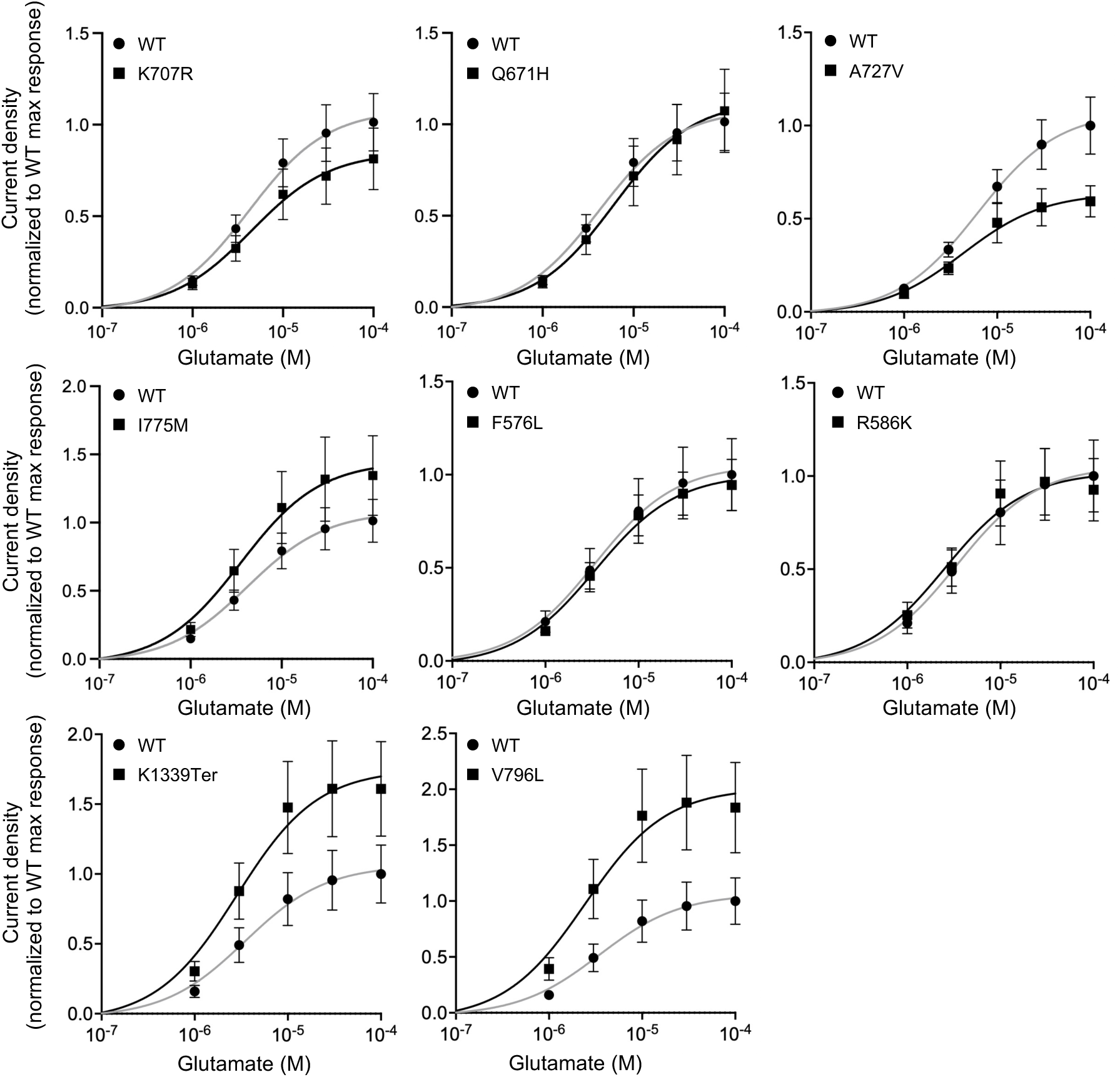
Effect of non-pathogenic *GRIN2A* variants on NMDAR function. Averaged current density in response to increasing concentrations of glutamate in the constant presence of 30 μM glycine, normalized to maximal response, for NMDARs containing the wild-type or non-pathogenic GluN2A variants. The lines indicate a nonlinear regression three-parameter fit to each dataset. Data are displayed as mean ± SEM, n = 17-70; see Table 1 for number of cells recorded per variant.

**Figure S5.**
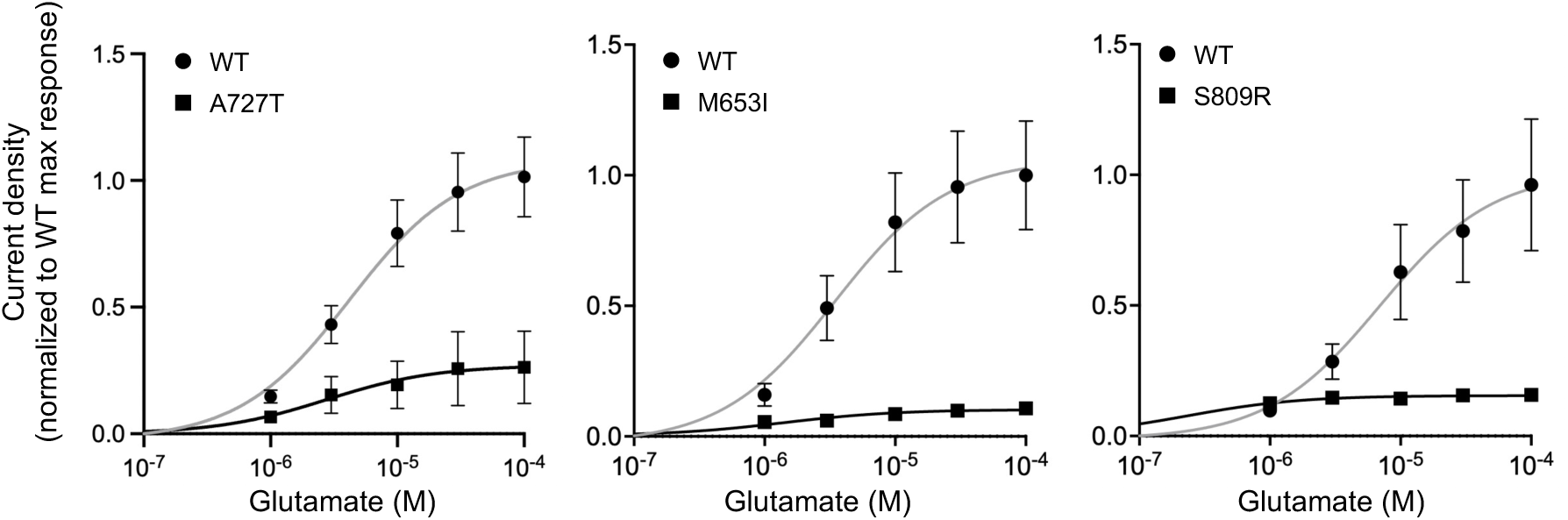
Effect of Epilepsy and DD/ID-associated *GRIN2A* variants on NMDAR function. Averaged current density in response to increasing concentrations of glutamate in the constant presence of 30 μM glycine, normalized to maximal response, for NMDARs containing the wild-type or Epilepsy and DD/ID-linked GluN2A variants. The lines indicate a nonlinear regression three-parameter fit to each dataset. Data are displayed as mean ± SEM, n = 10-77; see Table 1 for number of cells recorded per variant.

**Figure S6.**
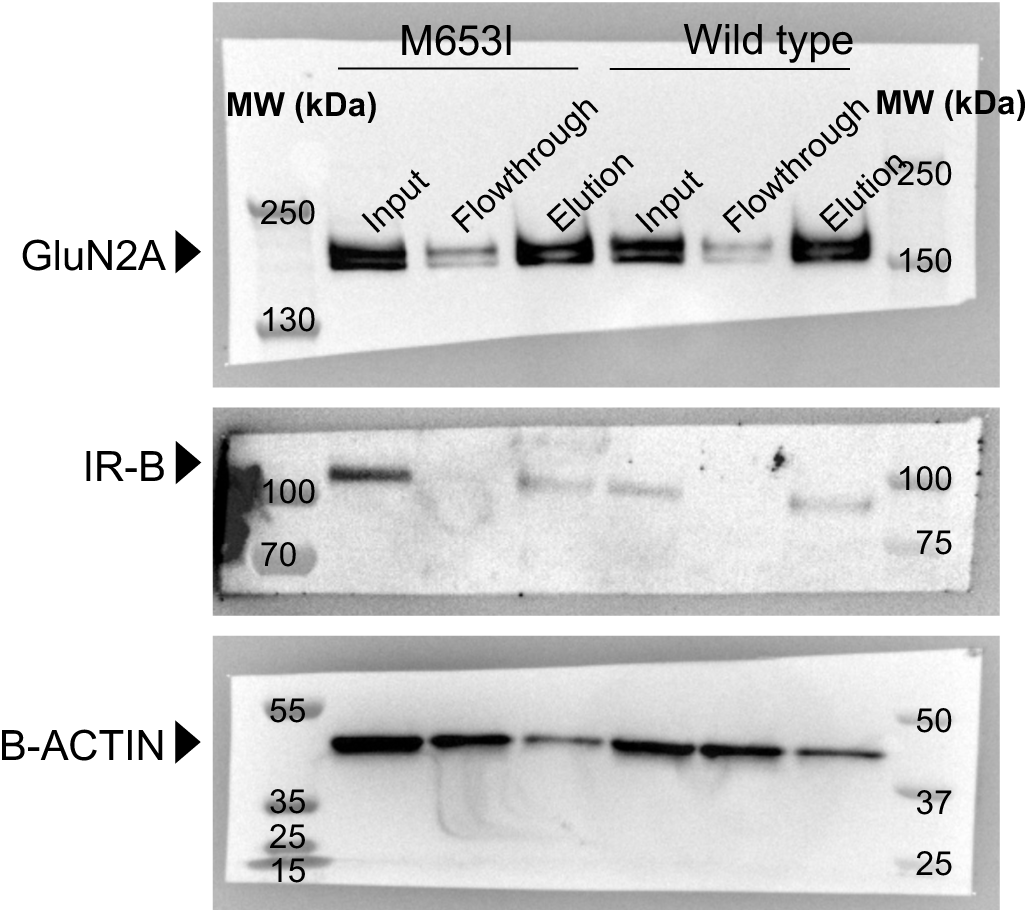
Original western blot images of the surface biotinylation experiment. Western blot probing for GluN2A, insulin receptor beta, and β-ACTIN in the input, flowthrough, and elution samples of surface biotinylation experiment done on HEK cells transiently transfected with *GRIN1-GRIN2A* constructs to express wild-type or M653I mutant NMDARs. Input, flowthrough, and elution represent total, internal, and surface expression respectively. IR-B: insulin receptor beta.

